# Preserved social development but impaired executive function in a *Shank3*-deficient rat model of Phelan-McDermid syndrome

**DOI:** 10.64898/2026.08.13.744651

**Authors:** Anna C. Pearson, Taylor M. Drazan, Sean P. Bradley, Audrey Thurm, Joseph D. Buxbaum, Jill L. Silverman, Yogita Chudasama

**Author notes:** Address for correspondence: Yogita Chudasama, Ph.D. National Institute of Mental Health Building 35A, Room 2D-804 Bethesda, MD 20892.

## Abstract

Phelan-McDermid syndrome (PMS) is a genetic neurodevelopmental disorder caused by a microdeletion within chromosome 22q13.3 ^1–6^, and accounts for ∼1-3% of cases of autism spectrum disorders (ASD). Individuals with PMS typically present with neonatal hypotonia, severe speech delay, intellectual disability, motor impairments, and autism-related features, although additional manifestations such as epilepsy, sleep disturbances, and developmental regression are common. As in ASD, there is considerable heterogeneity in cognitive and behavioral impairments in PMS, making it difficult to determine which features arise directly from *SHANK3/Shank3* deficiency and how deficits manifest across development. In the present study, we transferred the targeted *Shank3* mutation, previously characterized on the Sprague Dawley background, onto the Long-Evans strain and performed a longitudinal behavioral assessment of *Shank3*-deficient rats from infancy to adulthood. The rats were evaluated for delays in early sensory and motor development, and ultrasonic vocalizations as neonates, social behavior, short term memory and gait as juveniles, and cognitive-executive dysfunction using a touchscreen-based visual discrimination and reversal learning task as adults. Despite profound reductions and/or complete loss of *Shank3* protein expression, heterozygous and homozygous male and female rats showed normal physical and neurological reflexes across development, yet female *Shank3* knockout pups showed a selective reduction in distress-associated ultrasonic vocalizations. As juveniles, subtle abnormalities emerged in short-term memory and limb coordination, while social preference for novelty and recognition was normal. Prominent behavioral abnormalities were observed in adults characterized by rapid and error-prone responding to visual stimuli during discrimination learning and reversal. These data suggest that rats with complete or partial *Shank3* deficiency produce a selective behavioral profile in which high-order cognitive dysfunction is more pronounced than deficits in basic sensorimotor or select social behaviors. More broadly, these findings highlight executive dysfunction as a key consequence of the loss of *Shank3*. For the first time, we report the loss of *Shank3* expression on a Long-Evans background strain as a valuable translational tool for investigating cognitive dysfunction and therapeutic windows in *Shank3*-associated disorders.

## INTRODUCTION

*SHANK3* encodes a postsynaptic scaffolding protein essential for glutamatergic synapse development and function^1,2^. A deletion or mutation of *SHANK3* on chromosome 22q13.3 causes Phelan-McDermid Syndrome (PMS), a rare genetic neurodevelopmental disorder characterized by intellectual disability, language impairment, motor dysfunction and variable levels of autism-related features^3–5^.Clinical manifestations of PMS exhibit substantial variability in cognitive and behavioral outcomes^6^. Increasing evidence indicates that deficits in executive function, particularly inattention and cognitive inflexibility, are a prominent feature of PMS and may be especially pronounced even in individuals with point mutations or small deletions^7^. These impairments are thought to contribute substantially to difficulties with learning and goal-directed behavior underscoring the importance of executive dysfunction as a translationally relevant phenotype.

Several mouse models have advanced our understanding of the molecular and synaptic mechanisms of *Shank3* loss^2,8–10^, but have been less successful in modeling the higher-order cognitive manifestations associated with PMS^11^. This likely reflects, in part, the relatively limited cognitive repertoire of mice and the considerable variability in behavioral phenotypes across different genetic constructs and testing paradigms. The original *Shank3*-deficient rat model extended this work to a species with greater cognitive and social complexity, and has revealed impairments in attentional control and social communication^12,13^. However, characterization of this model has been exclusively restricted to the Sprague-Dawley strain. Compared with Sprague-Dawley rats, Long-Evans rats exhibit better visual function, greater exploratory behavior, and enhanced performance in cognitively demanding tasks, making them particularly well suited for studies of learning, attention, and executive function^14,15^. These differences are especially relevant for touchscreen-based assessments because Sprague-Dawley rats are albino and associated with altered retinal organization and reduced visual acuity^20–23^. Since PMS is a neurodevelopmental disorder and *Shank3* expression peaks postnatally in cortico-striatal brain structures^16–18^, understanding when behavioral abnormalities first emerge is as important as identifying their adult manifestations. This issue is critical for translational research, as it is particularly relevant for higher-order cognitive functions which are commonly impaired in PMS but difficult to assess systematically in affected individuals because of intellectual disability, poor motor skills and language limitations^19,20^. To address this knowledge gap, we performed a longitudinal assessment of *Shank3*-deficient rats generated and maintained on a Long-Evans background strain, from infancy to adulthood to define the developmental emergence of social, emotional, motor, and cognitive abnormalities relevant to PMS.

## METHODS

### Subjects

Male and female *Shank3* Sprague Dawley heterozygous rats were obtained as a gift from Dr. Joseph Buxbaum at Mount Sinai School of Medicine and rederived at the National Institute of Mental Health (NIMH) Transgenic Core facility on an outbred Long-Evans background. Full details of the generation of the *Shank3* gene mutation are provided in Harony-Nicolas et al (2017)^13^. Heterozygous breeder rats were paired to generate the animals used in this study and backcrossed for at least seven generations (**Fig. 1A**). We used male and female Long-Evans rats that were wildtype for *Shank3*^+/+^ (WT, n = 16), heterozygous for *Shank3*^+/-^and *Shank3*^-/+^ (HET, n = 34), and homozygous knockouts for *Shank3*^-/-^(KO, n = 18). These rats were tested as neonates and thereafter as juveniles and adults in a single cohort.

**Figure 1.**
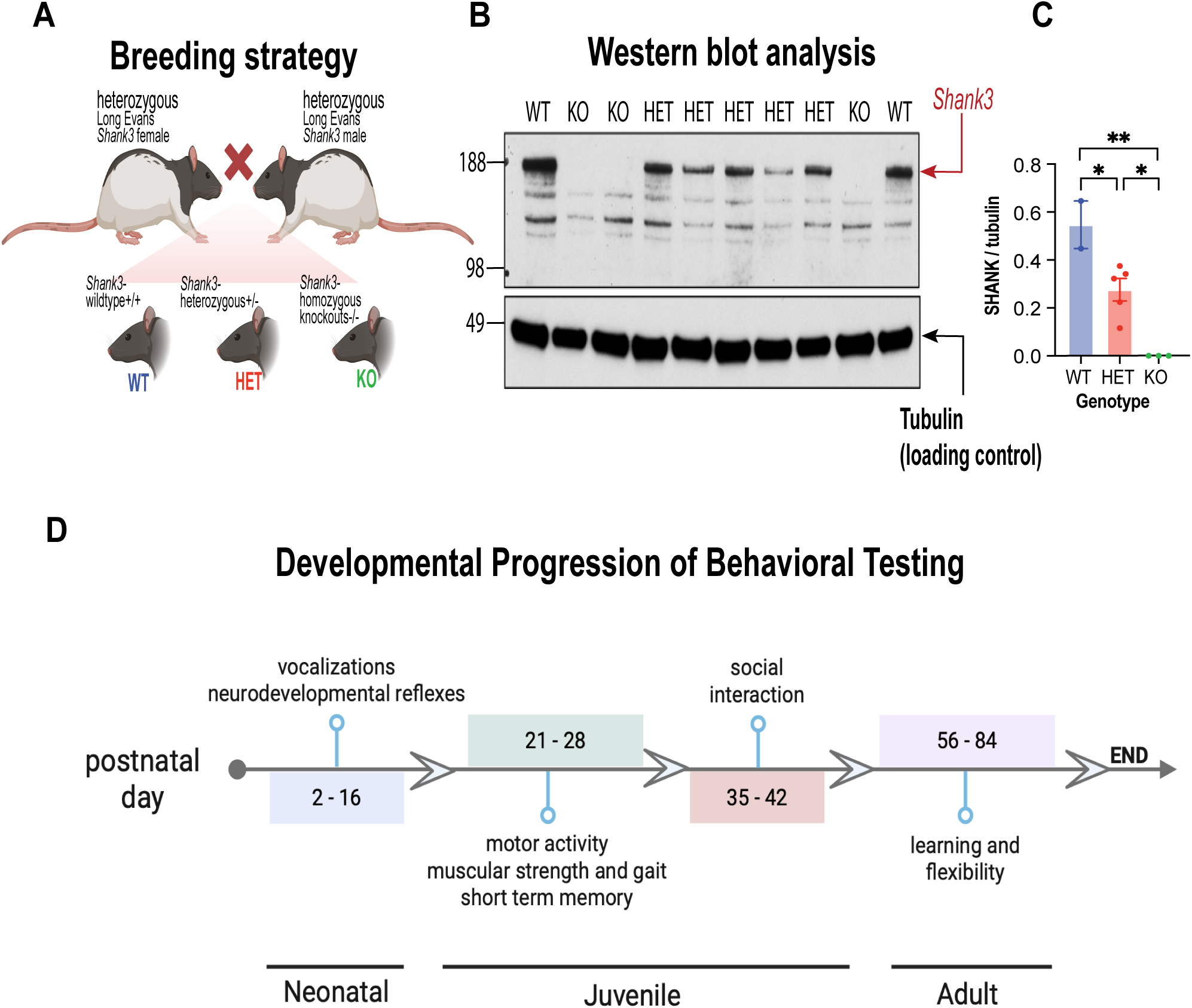
Validation and longitudinal assessment of Shank3 mutant Long-Evans rats. **(A)** Breeding strategy used to generate experimental animals. Heterozygous Long-Evans Shank3 mutant females were crossed with heterozygous Long-Evans Shank3 mutant males to produce wildtype (WT), heterozygous (HET), and homozygous knockout (KO) offspring https://BioRender.com/xu25j0e. **(B)** Representative Western blot showing Shank3 protein expression across genotypes. Shank3 immunoreactivity was detected at approximately 180 kDa, and tubulin was used as a loading control. **(C)** Quantification of Shank3 protein expression normalized to tubulin. Shank3 expression was reduced in HET animals and absent in KO animals relative to WT controls. Data points represent individual animals. Bars indicate mean ± SEM. ** p < 0.01; * p < 0.05. **(D)** Experimental timeline to show longitudinal behavioral assessment of Shank3-deficient rats from infancy to adulthood.

Behavioral testing was scheduled to minimize fatigue, repeated-testing effects, and potential carryover between assays while preserving the longitudinal design^21,22^. The order of testing was held constant across animals and was arranged to progress from short procedures to more socially or cognitive demanding assays as adults. A minimum of 24 hours separated successive tests. Ultrasonic vocalizations (USVs) were recorded on PD7 and PD14. Juvenile assessment proceeded in the following order: open field, gait analysis, grip strength, and novel object recognition (NOR). Social interaction testing was conducted at least 7 days after NOR, and touchscreen training began at least 2 weeks after the completion of social testing. All behavioral testing was conducted at approximately the same time of day to reduce variability associated with circadian influences on activity and performance.

Upon weaning at postnatal day (PD) 21, rats were group-housed with mixed genotypes in a temperature-controlled room (23.3°C) with a 12:12h light:dark cycle according to ARRIVE guidelines^23^. All behavioral testing occurred during the light phase. All experimental procedures were approved by the NIMH Institutional Animal Care and Use Committee, in accordance with the NIH guidelines for the use of animals.

### Genotyping and Western blotting for DNA, and protein expression

On PD1-3, pups were tattooed with a non-toxic marker for identification. Tail clippings for genotyping took place at PD21. Thus, all neonatal testing (PD2-16) was conducted blind to the genotype and before tail clippings so as not to affect their neonatal milestones including vocalizations. Genotypes were confirmed with qPCR for RNA through TransnetYX Inc (Cordova, TN). In addition, to confirm the reduced expression of *Shank3* in the *Shank3*-deficient rats, we performed a Western Blot analysis. Hemibrain lysates were prepared as outlined in Bozdagi et al. (2010)^2^, washed in a 4-12% Bis Tris 12-Well Gel (Invitrogen, #NPO322BO) and NuPAGE™ MES SDS Running Buffer (20X, Invitrogen, #NP0002), and transferred a polyvinylidene fluoride membrane. Anti-Shank 3 (1:1000, Cell Signaling Technologies, DK56R, 64555S Rabbit) and anti-beta III tubulin (1:3000, Sigma, T9026 Mouse) antibodies were used for the detection of Shank 3 protein and beta III tubulin, respectively. SeeBlue™ Plus2 Pre-Stained Protein Standard was used as a marking ladder (Invitrogen, #LC5925).

### Behavioral testing

#### Neonatal rats (postnatal day 2 – 16)

##### Neurodevelopmental reflexes

To capture physical and neurological reflex delays and potential symptoms of hypotonia (low or weak muscle tone) in early life, we implemented a series of short behavioral tests adapted from Fox (1965)^24^ to newborn rats from PD2 to PD16^25^. We focused on tests that assessed neurological reflexes which are involuntary and repetitive movements that demonstrate brain stem and spinal cord function (**Fig 2A, B**). In brief, we examined: 1) body rotation and coordination (surface righting and negative geotaxis), 2) strength and vestibular imbalance (forelimb grasp and cliff aversion), and 3) sensory system maturation (rooting, auditory startle, ear twitch). A description of each test can be found in supplementary materials (**Table S1**).

**Figure 2.**
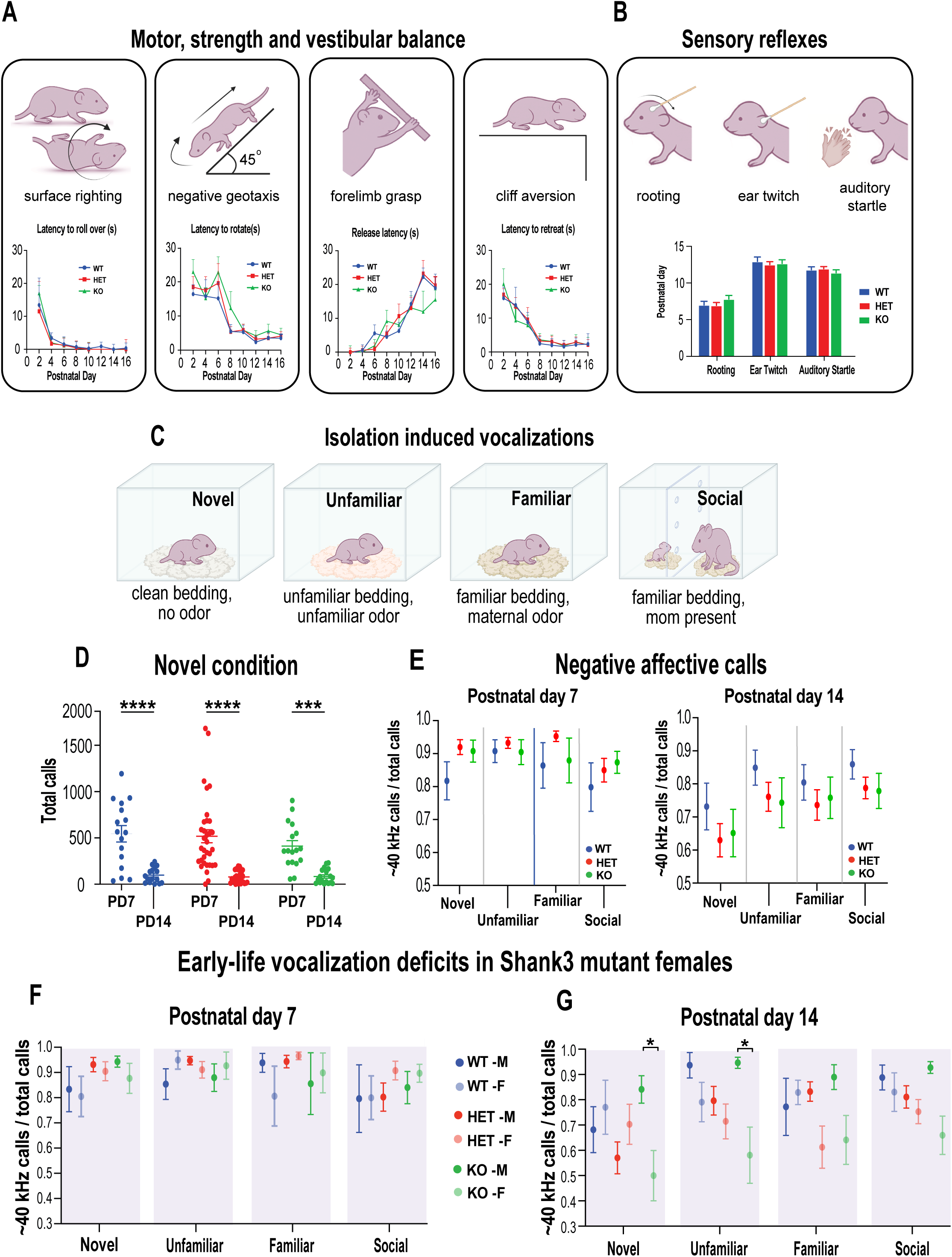
Early sensory and motor development is preserved, but affective vocalizations are altered in female Shank3 deficient pups. **(A)** Assessment of neonatal motor strength, and vestibular reflex development from postnatal day (PD) 2-16. Pups were tested for surface righting (latency to roll from a supine to prone position), negative geotaxis (latency to rotate 180° on a 45° inclined plane), forelimb grasp (release latency from a suspended rod), and cliff aversion (latency to retreat from a platform edge). https://BioRender.com/355e0eb. Performance improved in all animals with no significant genotype differences. Data are presented as mean ± SEM. **(B)** Development of sensory reflexes. The age of onset for rooting, ear twitch, and auditory startle reflexes were recorded. Reflexes emerged at comparable ages across genotypes indicating normal sensory development in Shank3 mutant rats. **(C)** Schematic of experimental conditions used to assess isolation-induced ultrasonic vocalization (USVs) https://BioRender.com/4vq9wf5. Pups were recorded under four conditions varying in familiarity and social context: Novel (clean bedding, no odor), Unfamiliar (unfamiliar bedding and odor), Familiar (home-cage bedding containing maternal odor), and Social (familiar bedding with the dam present). **(D)** Total USV production in the Novel condition at PD7 and PD14. Call rates declined with age across all genotypes, consistent with normal developmental reduction of isolation-induced USVs. \*\*\**P* < 0.001, \*\*\*\**P* < 0.0001. **(E)** Proportion of negative affective calls (∼40-kHz) relative to total calls at PD7 and PD14. These calls were largely preserved across conditions and genotype. **(F–G)** Sex-specific analysis of negative affective calls. **(F)** At PD7, the proportion of distress-associated calls did not differ between males and females in any genotype. **(G)** At PD14, female Shank3 knockouts (KO) pups emitted a lower proportion of negative affective calls relative to WT controls, with the most pronounced differences observed in teh novel and unfamiliar isolation conditions indicating altered affective vocal communication. *P < 0.05. Data are presented as mean ± SEM; M, male; F, female.

##### Vocalizations

Isolation induced ultrasonic vocalizations (USVs) were recorded in pups on PD7 and PD14 **(Fig. 2C**) to capture developmental changes in vocal behavior across a period when isolation-induced USVs undergo marked changes in call rate, frequency, duration and acoustic structure during the first postnatal weeks^26–28^. At both timepoints, vocalizations were assessed under four sequential conditions to assess whether affective vocal output varied as a function of environmental and social context. This allowed us to compare vocalizations across a graded set of conditions ranging from minimal stimulation in isolation to increasing levels of familiarity and social relevance. The design allowed us to determine not only whether *Shank3-*deficiency altered the normal developmental trajectory of pup vocalizations, but also whether vocal responses were appropriately modulated by changing environmental and social context.

Pups were separated from their mothers and placed individually on a heat pad in a Plexiglass recording chamber (L 41 cm × W 20 cm × H 20 cm). Pup vocalizations were recorded for three minutes in each condition. The recording conditions were run in the following order: 1) Novel, pup was alone in chamber with clean cage bedding; 2) Unfamiliar, pup was alone in chamber with cage bedding from an unfamiliar virgin female; 3) Familiar, pup was alone in chamber with cage bedding from its home cage; and 4) Social Interaction, the pup’s mother was placed in testing chamber behind a clear Plexigass wall with ventilation holes. In between conditions, the pup was placed on a heat pad in a padded Plexiglass holding chamber while the recording chamber was cleaned with 70% ethanol and allowed to air dry for three minutes before the next condition. Immediately after the final recording, pups were returned to their littermates. Litters were reintroduced to the mother after all recordings were complete. The entire procedure took 25 minutes.

Vocalizations were acquired using an Avisoft CM16/CMPA condenser microphone and connected via an UltraSoundGate 416H USB audio device to a computer with Avisoft-RECORDER software (Avisoft Bioacoustics, Berlin, Germany). The recorded sounds were digitized at a sampling frequency of 250 kHz and analyzed with RavenPro (Cornell Lab of Ornithology; Ithaca, NY). Calls were identified by two band-limited entropy detectors, each with an entropy above background threshold of 1.5 dB. Calls were classified as 40kHz-like when captured by a detector band-limited from 30 kHz to 50 kHz. Calls were classified as 66kHz-like when captured between 55 kHz and 75 kHz. To avoid double-counting or harmonics, in cases where calls overlapped in both the 40 kHz and 66 kHz bands, only the putative vocalization with the highest entropy was counted.

#### Juvenile rats (postnatal day 21 – 42)

##### Motor control and coordination

To evaluate body movement, posture and gait, juvenile rats were placed in a clear Plexiglass square arena (L 40 cm × W 40 cm × H 29.8 cm) for five minutes in red light (i.e., darkness). Footprints were detected and recorded with a high-speed digital Basler Ace2 camera (Ahrensberg, Germany) located beneath the clear arena. Gait features were analyzed with FreeWalkScanTM2.0 tracking software (Clever Systems, Inc., Virginia, USA). Distance traveled was determined using TopScan (Clever Systems, Inc., Virginia, USA).

##### Muscular strength

A Bioseb (BIO-GS4) grip strength monitor was used to assess muscular strength. The animal was allowed to grasp a metal grid while gently pulled backwards in a horizontal plane to determine the maximal peak force. The highest force exerted within three attempts was recorded.

##### Open field

To assess locomotor activity, juvenile rats were placed in a square arena (L 44 cm× W 44 cm × H 40 cm) for thirty minutes. Animal movement was recorded with a digital camera positioned above the arena. Distance traveled and velocity were quantified using TopScan tracking software (Clever Systems, Inc., Virginia, USA). The arena was cleaned with 70% ethanol between trials to remove olfactory cues.

##### Novel object recognition

We next assessed the rats’ ability to recognize and remember previously encountered objects as a measure of short-term recognition memory. Animals were placed in the same arena used for the open field containing two identical objects. The two objects were falcon tubes (H 12.0 cm × 1.7 cm diameter) secured with glue on a gray plaque and placed in opposite corners of the arena. Rats were allowed to freely explore the objects for five minutes during the acquisition phase. The objects were validated during the acquisition phase by showing no baseline preference for either object. Animals were then returned to their home cage for a retention interval of 60–90 minutes. During the test phase, rats were returned to the arena in which one familiar object was replaced with a novel object of different shape and texture (lego construction, H 12.8 cm × W 6.4 cm × L 3.84 cm) and allowed to explore the objects for five minutes while behavior was recorded with an overhead camera. Object exploration was defined as directing the nose toward the object within approximately 1 cm and/or touching the object with the nose or forepaws. The left/right location of the novel object was counterbalanced across animals to avoid object or side biases. The arena and objects were cleaned with 70% ethanol between trials to remove olfactory cues.

##### Social interaction

General sociability was assessed using the standard three chambered social interaction test according to previously described protocols^29^. Animals were tested at postnatal day 35-42 (5-6 weeks). The Plexiglass chamber (L 44.5 cm × W 106.7 cm × H 45.7 cm) was divided into three chambers (left, right, and center) separated by dividers with arched openings to allow rats free and easy access to any compartment **(Fig. 3A).** On day 1 (habituation), test rats were acclimated to the empty apparatus for 10 minutes and allowed to explore two holding chambers (H 22 cm x W 12 cm x L 16 cm), one located in each distal chamber. On day 2 (social preference), a novel, unfamiliar stimulus rat was placed under one holding cup (left or right chamber) while the other cup remained empty. The test rat was placed in the center chamber and allowed to freely explore all three chambers for 5 minutes. On day 3 (social recognition), a second novel stimulus rat was placed under a holding cup in one chamber while the other holding cup contained the stimulus rat from day 2, which now served as a familiar rat. Again, the test rat was placed in the center chamber and allowed to explore both animals in each chamber. The stimulus rats were age-, strain-, and sex-matched to the test rat but were housed in a separate room in the vivarium to ensure novelty. The left/right side on which the stimulus rat was placed alternated between test subjects. Automated video tracking with TopScan (Clever Systems, Inc., Virginia, USA) was used to determine time spent in each zone.

**Figure 3.**
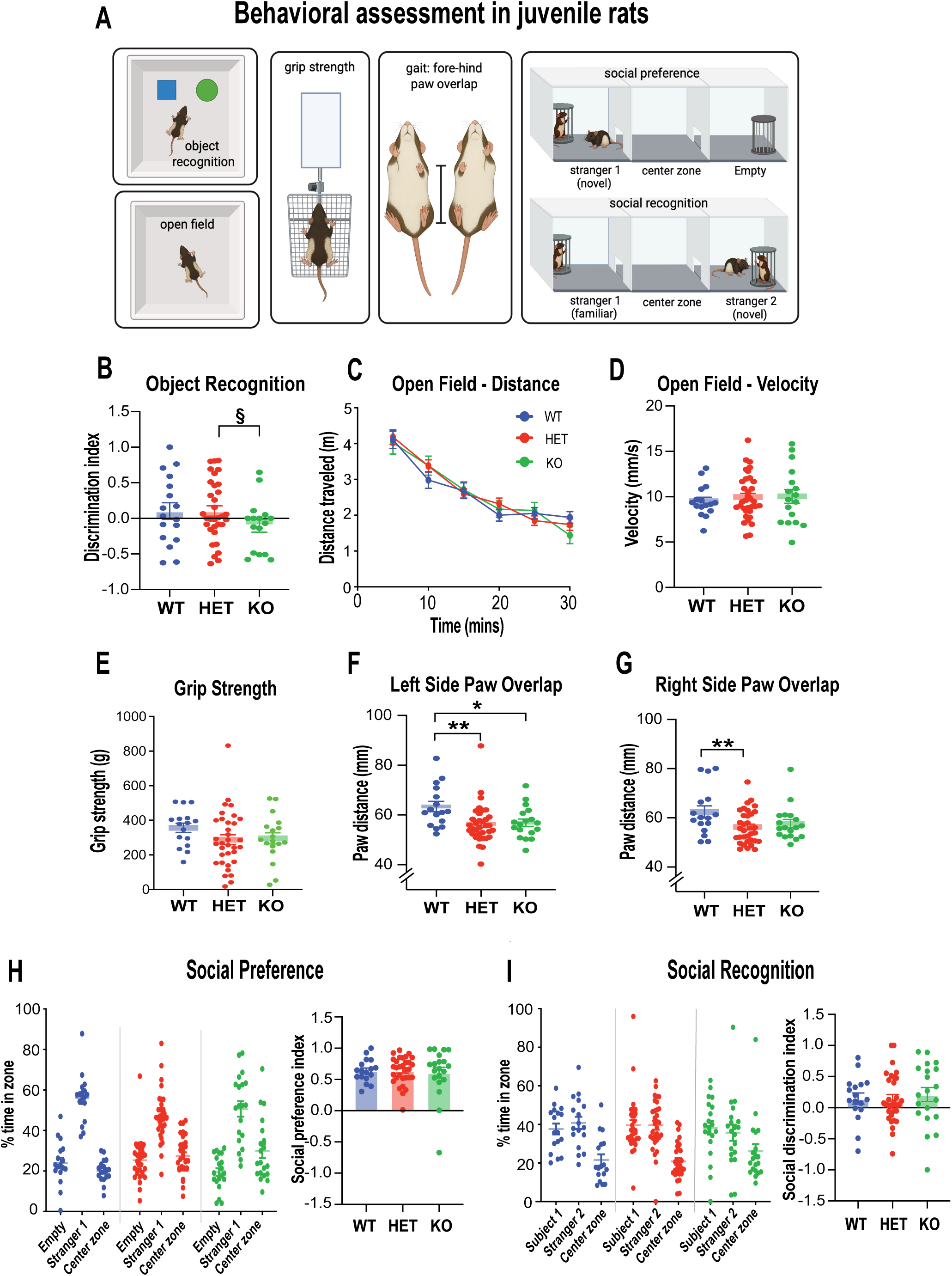
Juvenile Shank3-deficient rats exhibit subtle memory and motor coordination deficits but intact social behavior. **(A)** Schematic overview of behavioral assessments conducted in juvenile rats (PD21-PD28) https://BioRender.com/xhch7i9. **(B)** Novel object recognition performance expressed as a discrimination index. Shank3 knockout (KO) rats showed a trend toward reduced discrimination compared with heterozygous (HET) rats with only two rats above the discrimination threshold. § P = 0.07. **(C-D)** Open-field locomotor activity. (C) Total distance traveled during the 30-min session, and **(D)** average locomotor velocity did not differ among genotypes. **(E)** Forelimb grip strength was comparable across groups indicating preserved muscular strength. **(F–G)** Gait analysis of ipsilateral fore–hind paw coordination. **(F)** Left-side paw overlap was reduced in both HET and KO rats relative to WT controls. **(G)** Right-side paw overlap was reduced in HET rats whereas KO animals did not differ from either group. **(H)** Social preference test. Juvenile rats spent more time in the zone containing an unfamiliar conspecific (Stranger 1) than in the empty chamber regardless of genotype, indicating intact sociability. The social preference index (right panel) did not differ among groups. **(I)** Social recognition test. Following exposure to Stranger 1 during social preference, animals were tested with the familiar rat (Stranger 1) and a novel conspecific (Stranger 2) for social recognition. Investigation times and the social discrimination index (right panel) were comparable across genotypes, indicating intact social recognition memory. In all graphs for this figure, data are presented as mean ± SEM. WT, wildtype, HET, heterozygous; KO, homozygous knockout. Data points represent individual animals. *P < 0.05; **P < 0.01.

##### Adult rats (postnatal day 56 and above)

Cognitive impairments associated with the *Shank3*-deficiency (WT, n=10; HET, n=11; KO, n=17) were assessed in rats during performance of a visual discrimination and reversal learning task conducted in automated operant touchscreen chambers. For this task, adult rats were maintained at 85% of their free-feeding weight and had access to water for at least one hour each day. The apparatus and online data collection were controlled by Animal Behavior Environment (ABET) II software (Lafayette Instruments, Indiana, USA). Each chamber was individually housed within a sound-attenuating cabinet, ventilated by low-level noise fans, and illuminated by a 3W house light mounted on the ceiling of the sound-attenuating cabinet. The ceiling and two sidewalls of the operant chamber were made of clear Plexiglas (one of which served as the door). One side of the operant chamber was fitted with a touch-sensitive monitor (9”W x 10”H). Images of white squares were presented at two of five discreet locations on the touchscreen (**Fig. 4**). A black mask, approximately 1 cm from the surface of the display, served to restrict the rat’s access to the display except through response windows each measuring 2.06”L x 2.06”H. Opposite the touchscreen, on the rear wall, was a food magazine attached to liquid dispenser which delivered 10% sucrose as food reward.

**Figure 4.**
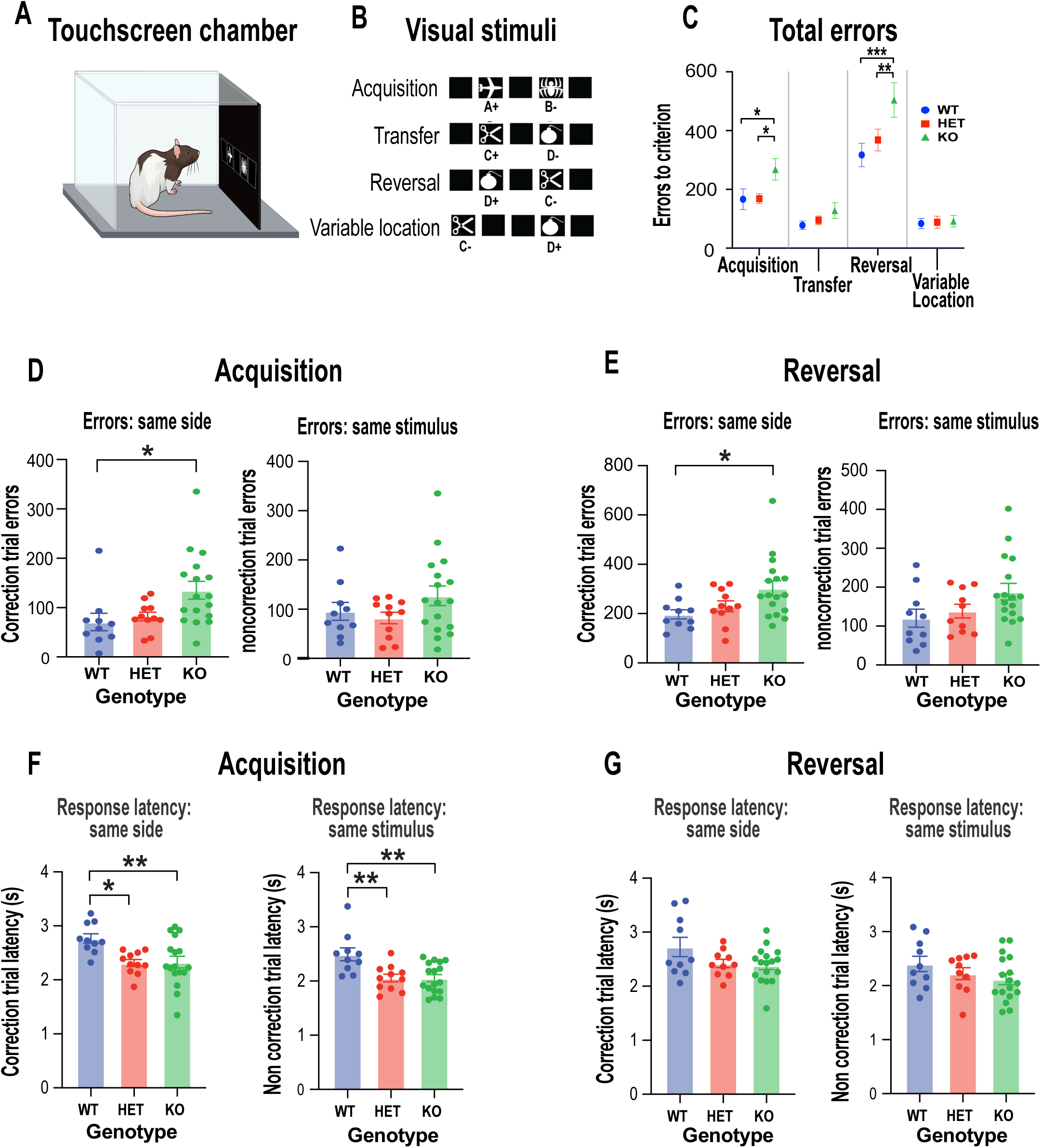
Shank3 deficiency impairs visual discrimination learning and cognitive flexibility through increased perseveration. **(A)** Schematic of operant touchscreen apparatus used for visual discrimination and reversal learning in adult rats. https://BioRender.com/u86mxln. **(B)** Visual stimuli and task structure. Animals first acquired a visual discrimination between two stimuli (A+ and B−), transferred the learned rule to a novel stimulus pair (C+ and D−), adapted to a reversal of reward contingencies (D+ and C−), and finally completed the discrimination with stimuli presented in variable spatial locations. **(C)** Total errors required to reach criterion during each phase of testing. Shank3 KO rats committed more errors than WT and HET animals for both acquisition and reversal. **(D)** Error analysis during acquisition. The KO rats made more correction-trial errors resulting in repeated responses to the same side (left panel) but same-stimulus non-correction trial errors did not differ between genotypes (right panel). **(E)** Error analysis during reversal learning. The KO rats exhibited an increase in same-side correction errors relative to WT animals (left panel), while same-stimulus non-correction errors remained unaffected (right panel). **(F)** Response latencies during acquisition. Both HET and KO rats responded more rapidly than WT controls on both same-side correction trials (left panel) and same-stimulus non-correction trials (right panel), indicating a faster but less accurate response strategy during initial learning. **(G)** Response latencies during reversal learning. No genotype differences were observed for either same-side correction-trials (left panel), or same-stimulus non-correction trials (right panel). In all cases, data are presented as mean ± SEM., with data points representing individual animals. WT, wild-type, HET, heterozygous; KO, homozygous knockout. *P < 0.05; **P < 0.01.

One week before testing, rats were given free access to dishes containing 10% sucrose in their home cage for 1-2 hours. Rats were periodically observed to ensure they were drinking the sucrose. Next, rats were acclimated to the touchscreen chamber for two 20-minute sessions and given free access to a small amount of 10% sucrose in the liquid receptacle. When rats were reliably consuming all the liquid in the receptacle, they were trained to make a nose poke touch to initiate trials. Initially, all five stimulus positions were presented, and a nose poke to any stimulus resulted in the delivery of a 50 µl sucrose reward. Once rats were reliably touching the screen and receiving 50 rewards within 20 minutes (across two sessions), the white squares were restricted to two positions (positions 2 and 4). A touch to either of these two stimuli within 10 seconds was rewarded with 50 µl of sucrose. The next trial was initiated when the rat made a nose-poke entry into the food receptacle. When rats made 50 touches in 20 minutes, they were ready for the main test session.

During the visual discrimination task, two novel geometric computer graphic stimuli (A+/B-) were presented in positions 2 and 4 **(Fig. 2B**). The left or right position of each stimulus was determined pseudorandomly. These stimuli remained on the screen until the rat made a nose-poke touch response to either stimulus. A correct response to one stimulus (designated A+) was rewarded. An incorrect response to the other stimulus (designated B-) was not rewarded and instead resulted in the disappearance of both stimuli from the screen, and a 5 s timeout period during which all the lights were extinguished. An incorrect response to B-resulted in a correction trial in which the same trial was repeated (i.e., the A+ and B-stimuli remained in the same left/right positions) until the rat responded correctly. Thus, correction trial errors were repeat errors made to the same side, whereas non-correction trial errors were made to the same stimulus. Each session could have an infinite number of correction trials but was limited to a total of 60 non-correction trials. Criterion performance was set to 85% accuracy for individual rats on two consecutive sessions.

To establish if the animals’ ability to transfer the stimulus-reward discrimination rule, rats were introduced to a second pair of novel stimuli (C+ and D-), after which the stimulus-reward contingencies were reversed so that the previously non-rewarded stimulus (i.e., D-) became the rewarded stimulus (i.e., D+), and vice versa. The rat was now required to reverse its response by inhibiting its response to the previously rewarded stimulus and respond to the new rewarded stimulus. Criterion performance was 85% accuracy on two consecutive sessions. Finally, we assessed the animals’ ability to transfer the discrimination rule to stimuli located in variable spatial locations. That is, the stimulus pair C-and D+, were presented in any of the five spatial locations.

#### Data analysis

The behavioral data were analyzed using GraphPad Prism (version 11) and custom MATLAB (version R2025b) and RStudio (version 2025.09.1+401) scripts. Data were analyzed for sex differences. In the absence of sex differences, data from both male and female subjects were combined to form a single group. The developmental trajectory of each genotype was analyzed with a linear mixed model (LMM) to account for the variability in testing days. Where applicable, data were analyzed with one-, two-, or three-way ANOVA. The data for grip strength was analyzed using a Kruskal-Wallis test to account for unequal distributions.

## RESULTS

Pairing heterozygous rats successfully produced *Shank3*-wildtype^+/+^ (WT), *Shank3-*heterozygous^+/-^or heterozygous ^-/+^ (HET), and *Shank3*-homozygous knockouts^-/-^(KO) rats (**Fig. 1A**). In a subset of animals, we performed a western blot analysis to ensure that the litters from rederived rats onto a Long-Evans background were indeed *Shank3-*deficient. According to GenBank (NCBI:txid10116), the molecular weight of the rat *Shank3* protein is approximately 180 kDa. A densitometric analysis of Western blot bands was performed using Sciugo open-source software. Band intensities for *Shank3* were measured and normalized to the corresponding tubulin bands to control for protein loading. An intense band of size 188 kDa was observed in WT rats and a less intense band was observed for all the HET animals. In the *Shank3* KO, this band was completely absent **(Fig. 1B**). Relative protein expression levels were calculated as the ratio of Shank to tubulin for each sample. A one-way ANOVA with data were corrected with Tukey’s multiple comparisons test confirmed that the generated *Shank3* mutant Long-Evans rats were indeed lacking the *Shank3* gene (WT vs HET, p = 0.028; WT vs KO, p = 0.001; HET vs KO p = 0.014; **Fig. 1C**).

### *Shank3*-deficient pups show normal sensory and motor development

**Fig. 1D**. provides a timeline of the various tests conducted on the animals from neonatal to adult. We first assessed early life reflexes across PD 2-16 using a standardized neonatal test battery (**Fig. 2A, B**). Data were collected blinded to genotype. Since movement is restricted in pups for the first few weeks of life, we measured coordination and rotation as a proxy for motor development. Over the course of 16 days, all pups, regardless of genotype flipped onto their paws from a supine position (righting reflex: (F(2,2) = 0.135, p = 0.879) and quickly rotated their bodies 180 degrees to reposition themselves against gravity (negative geotaxis: (F_(2,2)_ = 8.564, p = 0.105; **Fig. 2A**). The negative geotaxis skill was slow at first but improved with increasing age (F_(7,2)_ = 56.034, p = 0.004). Grip strength was intact as well (F_(2,2)_ = 3.317, p = 0.232) and improved as the pups got older (F_(7,2)_ = 42.936, p < 0.001) indicative of normal neuromuscular development. There was also no evidence of visuospatial or sensorimotor impairment as assessed by the visual cliff task (F_(2,2)_ = 0.231, p = 0.805; **Fig. 2A**). Early life reflexes to measure sensory detection all developed normally (Rooting F_(2,65)_ = 0.98), Ear Twitch F_(2,65)_ = 0.82; Auditory Startle F_(2,65)_ = 0.62; all p > 0.05; **Fig. 2B**). Thus, despite the major deficiency in the Shank3 protein, the early development of sensory and motor function in both HETS and KO rats was completely intact.

### *Shank3* deficiency alters early life vocalizations in females

Ultrasonic vocalizations (USV) are elicited when a pup is separated from the dam or littermates. These isolation-induced calls serve as a distress signal and widely used as developmental indicators of nervous system maturation and/or emotional reactivity^30,31^. Since USV production changes rapidly during early postnatal weeks, the ages of PD7 and PD14 were selected to capture distinct stages in the ontogeny of isolation-induced vocalizations. Rat pup USVs are robust during the first postnatal week and subsequently decline as pups approach weaning, with PD14 coinciding with major sensory and behavioral transitions, including eye opening^26,32^. Unlike mice, isolation-induced vocalizations in rats can persist beyond eye opening and decline progressively over subsequent development^27,28^. Vocalizations were recorded under four different contexts ranging from novel to familiar (**Fig. 2C**).

Across all genotypes (WT, n=16; HET, n=34; KO, n=18), total USV production declined overall for all conditions including the novel condition between PD7 and PD14 (all p<0.01; **Fig. 2D; Table S2A,B**), consistent with the normal developmental reduction in isolation-induced vocalizations. An analysis of the call structure revealed that the proportion of negative affective calls (40kHz)^33,34^ remained high across all conditions and was not influenced by genotype at either age (PD7: F_(2, 63)_ = 2.650, p = 0.078; PD14: F_(2, 63)_ = 1.227, p = 0.300, **Fig. 2E; Table S2C,D**). While there was no effect of condition at PD7 (F_(2.725, 163.5)_ = 1.875, p = 0.141), an effect did emerge at PD14 (F_(2.672, 155.0)_ = 8.168, p < 0.0001, **Fig. 2E**). The most prominent genotype-dependent effects emerged in female *Shank3* mutants at PD14. While no sex-dependent differences were detected at PD7 (**Fig. 2F**), at PD14, *Shank3* KO females exhibited a reduced proportion of ∼40 kHz distress calls relative to KO males when tested in both the novel (Mean ± SEM: KO Males 0.841 ± 0.054, KO Females 0.500 ± 0.099, post hoc Tukey’s test, p = 0.025) and unfamiliar (Mean ± SEM: KO Males 0.946 ± 0.021, KO Females 0.580 ± 0.111, post hoc Tukey’s test, p = 0.012) isolation conditions (**Fig. 2G**). A similar differentiation between male and females was present in the familiar condition (Mean ± SEM: KO Males 0.889 ± 0.048, KO Females 0.641 ± 0.096) and during social interaction (Mean ± SEM: KO Males 0.927 ± 0.023, KO Females 0.659 ± 0.075) but the groups did not differ statistically (post hoc Tukey’s test for familiar, p = 0.236 and social interaction p = 0.147). Most call characteristics did not vary across genotypes. There was a modest increase in call entropy exhibited by the HET animals between PD7 and PD14 (post hoc Tukey’s test p = 0.012) whereas WT and KO animals showed no age-dependent change (**Fig. S1A**). Call duration and peak frequency were unaffected by genotype or age **(Fig. S1B,C; Table S2E,J)**. These data suggest that the loss of *Shank3* selectively alters the expression of affective vocalizations in female pups during a critical developmental period, particularly in contexts associated with novelty or environmental uncertainty.

### Subtle memory deficit emerges before adulthood in *Shank3* deficient rats

The novel object recognition test was conducted in juvenile animals (PD 21–28) as a short-term memory assay and to reveal subtle cognitive impairments that might emerge before adulthood. The discrimination index provided a quantitative measure of how well the animals distinguished between the familiar and novel objects. There was high inter-individual variability in discrimination performance with data points widely dispersed even amongst the WT controls. The complete KO animals tended to have lower discrimination indices compared to HETs (F_(2,63)_ = 2.124, p = 0.07; **Fig 3B),** suggesting a trend toward impaired recognition memory. Despite the wide range in individual scores, the *Shank3*-deficient rats tended to explore the novel object less. In fact, only two animals from this group showed memory for the novel object with high discrimination indices; the remaining 14 animals performed near or considerably below chance level. This effect was unrelated to general locomotor activity since their ambulatory movement in the open field was normal (F_(10, 384)_ = 0.7244, p = 0.70; **Fig. 3C**) as was their speed (F_(2, 64)_ = 2.648, p = 0.08; **Fig. 3D**).

### *Shank3* deficient juveniles show altered limb coordination

Developmental regression of motor skills is a key feature in patients with PMS^35,36^. We therefore examined motor function in animals lacking the *Shank3* gene as juveniles since these animals showed normal motor coordination as neonates. First, we measured muscular strength but there was no detectable deficit; all animals regardless of genotype were able to exert the typical force for its size and age during a test of grip strength (Kruskal-Wallis H = 3.68, p > 0.05; **Fig. 3E**). However, gait analysis revealed subtle coordination abnormalities. Ipsilateral paw overlap, which reflects the spatial and temporal coordination between forelimb and hindlimb movements on the same body side, were altered in a genotype-and side-specific manner. On the left side, juvenile rats both heterozygous for the *Shank3* mutation (Mean ± SEM: 1.36 ± 56.43 mm) and the complete KO rats (Mean ± SEM: 56.91 ± 1.50 mm) exhibited reduced fore–hind paw distance relative to WT controls (Mean ± SEM: 63.39 ± 2.10 mm (P < 0.01 corrected with Tukey’s multiple correction), with no difference between mutant groups (**Fig. 3F)**. On the right side, only HET rats showed a significant reduction in paw overlap compared with WT (Mean ± SEM: HET, 56.26 ± 1.15 mm; WT, 62.43 ± 2.51 mm, p = 0.03), whereas KO animals were comparable to the other groups (Mean ± SEM: 57.65 ± 1.67 mm). These findings suggest that the mutation influences limb coordination, with the effect appearing more pronounced on one side.

### Social behavior is intact in juveniles with *Shank3* mutation

Social deficits have been reported in *Shank3* deficient rodent models and are often considered a translationally relevant feature of *Shank3*-associated neurodevelopmental disorders^2,8^. We used the standard three-chamber assay to assess sociability and social novelty preference in juvenile rats, facilitating comparison with the extensive literature using this paradigm in mouse models of *Shank3*-deficiency. We found no evidence of impaired social behavior in *Shank3* deficient animals. All genotypes displayed intact social preference by spending more time in the zone containing the unfamiliar conspecific (Stranger 1) than the zone containing the empty cup (P < 0.05; **Fig. 3H, left panel**). Consistent with this finding, the social preference index, reflecting time spent in close proximity to Stranger 1 relative to the empty cup, did not differ by genotype (**Fig. 3H, right panel).** Likewise, when stranger 1 (now familiar) was paired with a novel conspecific (stranger 2), animals in all groups preferentially investigated the novel animal and neither investigation time in the novel versus familiar zones nor the corresponding discrimination index differed by genotype (P > 0.05; **Fig. 3I**). In addition, latencies to first investigate the familiar or stranger rat were comparable across groups **(Fig. S2)**. Thus, there was no evidence of this type of impaired social approach and recognition behaviors in juvenile *Shank3*-deficient rats, using this specific assay, as previously reported^12,13^.

### Side-specific perseveration in adult *Shank3* deficient rats

When rats reached adulthood (PD ≥ 60), we assessed cognitive performance in an operant touchscreen discrimination task requiring them to learn and update stimulus-reward associations. Animals were first required to learn a visual discrimination, transferred the learned rule to a novel stimulus pair, adapted their responses following reversal of the reward contingencies, and finally performed the discrimination when the stimuli when presented in variable spatial locations (**Fig. 4A, B).** Genotype had a major influence on task performance (F_(2, 138)_ = 7.368, P < 0.001, **Fig. 4C**). *Shank3* KO rats were impaired during the acquisition phase committing more errors to criterion than both HET and WT groups (Mean ± SEM: WT 166.4 ± 34.19; HET 153.7 ± 20.26; KO 257.6 ± 36.51). Despite this learning difficulty, the KO rats successfully transferred the rule to a novel stimulus pair performing comparably to WT and HET groups (p > 0.05). However, when the stimulus-reward contingencies were reversed, KO rats again were impaired, committing many errors relative to controls (Mean ± SEM: WT 317.2 ± 39.91, HET 368.2 ± 37.22, KO 472.4 ± 39.52). Once the reversed rule was learnt, all animals performed similarly during the variable-location phase (Mean ± SEM: WT 83.80 ± 17.32, HET 88.00 ± 20.83, KO 86.47 ± 19.10) indicating intact stimulus discrimination independent of stimulus position.

To better characterize the acquisition and reversal deficits, we examined patterns of errors on both correction and non-correction trials (see Methods). During acquisition, the KO animals committed more same-side errors than the WT rats (F_(2, 35)_ = 4.585, P = 0.017, **Fig 4D Left**), whereas same-stimulus errors did not differ across genotypes (F_(2, 35)_ = 1.747, P = 0.189, **Fig 4D Right**). A similar pattern emerged during reversal learning; there was an increase in same-side correction errors in the KO rats relative to WT (F_(2, 35)_ = 4.238, P = 0.022 **Fig. 4E Left**), but same-stimulus errors remained comparable across groups (F_(2, 34)_ = 2.871, P = 0.070, **Fig. 4E Right**). This would suggest that the *Shank3* deficient animals were driven by repeated responding to a previously selected spatial location rather than an inability to discriminate stimulus identity. Importantly, because the rewarded stimulus occurred equally often on the left and right side of the touchscreen, the increased same-side errors could not be attributed to a generalized side bias (for acquisition, F_(2, 35)_ = 0.6271, P = 0.540; for reversal, F_(2, 34)_ = 0.160, P = 0.852, **Fig. S3A, B**).

We further analyzed errors committed during performance on a trial-by-trial basis to establish the animals’ likelihood of making an error following consecutive incorrect trials (error runs). Across all genotypes, animals typically corrected an incorrect response within 2–3 trials and made progressively fewer errors following longer consecutive runs of errors. Although *Shank3* KO rats made more errors overall than WT and HET animals for both acquisition (F_(2,105)_ = 7.951, p = 0.0006) and reversal (F_(2,103)_ = 18.50, p < 0.0001), there was no genotype × error run interaction in either phase (all p > 0.05), indicating that the reduction in errors across successive error runs was similar across genotypes despite the overall impairment in KO animals **(Fig. S3C, D**). These data suggest that the increased error rate in *Shank3*-deficient rats was not attributable to prolonged perseverative sequences. Rather the impairment appears to reflect a greater overall tendency to repeat incorrect responses consistent with a difficulty disengaging from a previously executed response. This interpretation is supported by the short response latencies observed when the *Shank3* KOs repeated responses to the same side during acquisition (F_(2, 35)_ = 5.922, P = 0.006, **Fig. 4F Left**) and to the same stimulus (F_(2, 35)_ = 8.101, P = 0.001, **Fig. 4F Right)**. In contrast, response latencies during reversal and reward collection latencies were largely unaffected by genotype (**all P>0.05; Fig. G, Fig. S3E, F).**

## DISCUSSION

The present study provides the first longitudinal behavioral assessment of *Shank3*-deficiency in rats maintained on a Long-Evans background spanning neonatal, juvenile, and adult stages of development. The Long-Evans background offers important advantages for modeling higher-order cognitive features of PMS, including intact visual function and robust performance on cognitively demanding tasks, making it particularly well suited for touchscreen-based assessment of learning and executive function. Despite profound reduction or absence of Shank3 protein expression, mutant animals exhibited largely intact sensory and motor development, preserved social interactions, and modest abnormalities during the juvenile period. In contrast, the most prominent behavioral deficit emerged in adulthood during touchscreen-based cognitive testing, where *Shank3*-deficient rats displayed impairments in discrimination and reversal learning that were driven by repeated responding to previously selected locations. These findings establish the Long-Evans *Shank3*-deficient rat as a valuable model of PMS and reveal a developmentally evolving behavioral profile in which deficits in higher-order cognitive control are more pronounced than deficits in basic sensorimotor or specific social behaviors.

We observed several aspects of behavior that were unexpectedly preserved. Neither HET nor KO mutants exhibited delays in the acquisition of neonatal reflexes, sensory responsiveness, or gross motor development. Likewise, juvenile mutants showed a robust preference for a social stimulus over the nonsocial alternative with no evidence of impaired social recognition, findings that contrast with reports of social impairments of this type in some *Shank3* mouse models^2,8,37^. In rats, however, the absence of a preference for the novel conspecific should not necessarily be interpreted as impaired social recognition. Social novelty preference requires not only discrimination between familiar and novel individuals, but also a motivational bias toward investigating the novel animal. Rats are highly social and strongly motivated to engage in direct social contact, but because the three-chamber paradigm restricts direct physical interaction, continued investigation of the familiar conspecific may reflect persistent social motivation rather than a failure to recognize the familiar animal. Thus, the equivalent investigation of familiar and novel animals may reflect limitations in the expression of social recognition within the three-chamber paradigm rather than an inability to discriminate social familiarity. These species-specific considerations may be particularly important in Long-Evans rats and suggest that the social novelty phase of the three-chamber assay should be interpreted cautiously when used as a measure of social recognition in this strain. We note that previous studies of *Shank3*-deficient rats reported intact short-term social recognition despite impairments in more demanding forms of social memory^12,13,38^. Our KO rats showed a tendency toward impaired novel object recognition early in their juvenile period, suggesting that deficits in short-term memory may emerge before detectable impairments in social behavior. Although differences in genetic background and testing procedures may contribute to variation across studies, the preservation of aspects of sociability observed here mirrors the clinical heterogeneity seen in individuals with PMS, where social functioning varies substantially despite at least some shared genetic etiology^39^. Unlike rodent behavioral tasks that are designed to capture broad social deficits observed in other ASD-associated conditions, differences in social deficits associated with *Shank3* disruption may reflect specific attention and memory alterations that impact social functioning^40^

Although early sensory and motor development was largely preserved, several more subtle alterations emerged during early development. One subtle change was in affective vocal communication during infancy. Isolation-induced ultrasonic vocalizations are widely used as an index of emotional reactivity and changes in pup–dam communication are often interpreted as reflecting alterations in the neural circuits regulating affective state^12,30,31^. Consistent with normal development, overall call production declined substantially between PD7 and PD14 across all genotypes. However, by PD14, only female *Shank3* KO pups emitted a lower proportion of ∼40 kHz distress-associated calls, particularly when exposed to novel or unfamiliar environments. This effect was not present at PD7 and occurred in the absence of changes in total call output or call structure suggesting that *Shank3* deficiency selectively influenced the affective quality rather than the overall production of vocalizations. The female-specific nature of the effect suggests that the behavioral consequences of *Shank3* deficiency may, under some conditions, be modulated by sex-dependent developmental processes. However, sex differences are not a consistent clinical feature of PMS. Developmental behavioral studies in multiparous species like rats are inherently susceptible to litter-related variability, and subtle sex-specific effects require substantially larger sample sizes for robust confirmation. Nevertheless, the restriction of this effect to homozygous knockout animals suggests a more severe loss of *Shank3* function may be necessary for this alteration in affective vocal signaling to emerge.

A modest increase in call entropy was observed in HET animals during development. Entropy is commonly interpreted as a measure of acoustic complexity^28^, suggesting that distress calls became slightly less stereotyped with age in this group. However, the effect was restricted to heterozygous animals and was not accompanied by changes in call duration or frequency, indicating that it is unlikely to account for the more prominent female-specific alterations in affective vocal signaling observed in KO pups. Although robust sex differences in social functioning have not been consistently reported in individuals with PMS, both clinical and preclinical studies suggest that *Shank3*-related phenotypes may be differentially modulated in males and females^12,41^. Importantly, while the functional significance of the altered isolation induced distress-call profile remains unclear, these vocalizations have a communicative function in pup–dam interactions, as playback of pup isolation calls elicits maternal approach and retrieval behavior^12^. Thus, the selective reduction in distress associated calls may reflect a subtle alteration in early affective communication between pup and dam. Our findings further suggest that this behavior may be influenced by biological sex, and emerge early in life, even when overt behavioral impairments are not apparent.

*Shank3* deficiency did not disrupt the initial development of basic motor function, but more complex aspects of motor skill involving gait become affected when the rats were juvenile. This temporal dissociation is noteworthy given that motor regression and deterioration of motor skills are well-recognized features of PMS^35,36,42^. Although the present data do not demonstrate true regression, as motor performance was not tracked longitudinally within individual animals, the appearance of coordination deficits after a period of apparently normal development in infancy is consistent with the notion that *Shank3* deficiency may interfere with the maturation or maintenance of motor circuits rather than their initial formation. Such an interpretation is compatible with the established role of *Shank3* in synaptic stabilization and plasticity^43^ and suggests that motor abnormalities may emerge as neural circuits become increasingly refined during postnatal development. Future studies would need to determine whether these early coordination deficits remain stable or progress with age, to closely model the regressive motor features reported in some individuals with Phelan-McDermid syndrome.

The clearest behavioral abnormality in the KO rats was a deficit in learning and executive function observed in adulthood during touchscreen discrimination learning. *Shank3*-deficient rats required more trials to acquire and reverse stimulus–reward contingencies yet successfully transferred learned rules to novel stimuli and performed normally when stimulus locations varied. These deficits were not attributable to impaired stimulus discrimination, generalized side bias, or reduced sensitivity to negative feedback. Instead, the *Shank3* deleted animals repeatedly responded to previously selected response locations suggesting a difficulty disengaging from prior actions. A similar pattern of perseverative responding has been reported in *Shank3*-deficient mice in the Barnes Maze, where animals repeatedly returned to previously searched or previously reinforced escape locations despite changing task demands, supporting a conserved association between *Shank3* deficiency and impaired behavioral flexibility across species^44^. Intriguingly, the rats responded unusually fast during the acquisition phase; response latencies normalized during reversal and reward collection latencies remained intact throughout testing. Thus, during initial learning, *Shank3*-deficient rats were not slower or less engaged than controls but instead responded rapidly while making more errors. This pattern is consistent with an altered speed–accuracy tradeoff in which choices are executed hastily rather than engaging in deliberate stimulus evaluation. Together, these findings suggest that *Shank3* deficiency disrupts executive control over response selection, rather than learning capacity itself. This distinction is clinically important, as it suggests that *SHANK3* disruption may impair the ability to regulate and adapt behavior rather than producing a generalized cognitive deficit. Such a profile is consistent with dysfunction within corticostriatal networks that regulate action selection, cognitive flexibility, and inhibitory control^45,46^, systems that are known to be sensitive to *Shank3* disruption^9,47,48^.

In conclusion, we report that *Shank3* deficiency produces a selective and developmentally evolving behavioral profile in which higher-order cognitive dysfunction is more pronounced than deficits in basic sensorimotor or social behaviors. While several aspects of the phenotype were preserved, the evidence suggests that *Shank3* loss influences the maturation of neural circuits involved in affective communication, motor coordination, and short-term memory before the emergence of more pronounced cognitive deficits in adulthood. More broadly, the results mirror the marked heterogeneity observed in PMS and support the view that learning and executive dysfunction may represent a particularly sensitive consequence of *Shank3* disruption. By defining the developmental emergence of behavioral abnormalities, this work establishes the Shank3-deficient Long-Evans rat as a valuable translational model for investigating the neural mechanisms underlying cognitive dysfunction in *Shank3*-associated neurodevelopmental disorders and for identifying developmental windows that may be most amenable to therapeutic intervention.

## ACKNOWLEDGEMENTS

This research was supported by the Intramural Research Program of the National Institute of Mental Health (ZIA MH002951 and ZIC MH002952 to Y.C.), part of the National Institutes of Health (NIH). The contributions of the NIH author(s) are considered Works of the United States Government. The findings and conclusions presented in this paper are those of the author(s) and do not necessarily reflect the views of the NIH or the U.S. Department of Health and Human Services. Schematic illustrations of rats and behavioral assays were created with both Biorender.com and Adobe Illustrator v30.7; all other figure elements were generated by the authors. We thank Dr. Jim Pickle for rederiving the rats on a Long-Evans background and Mrs. Anamika Singh for training us in Western Blotting. We also thank the NIMH Rodent Behavioral Core staff for their help and guidance with behavioral testing infrastructure, testing protocols and training. We are indebted to the NIMH Veterinary Medicine and Resources Branch (VMRB) and NINDS Animal Health and Care Section (AHCS) for their services in animal health, hygiene and husbandry. ACP is currently a graduate student at Georgetown University, Washington DC, USA, and AT is now an Associate Professor at Boston Children’s Hospital in the Department of Psychiatry, Harvard Medical School, Boston MA, USA.

## SUPPLMENTARY MATERIAL

**Table S1.** Description and scoring of neurodevelopmental reflexes.

| Task | Description | Measure |
| --- | --- | --- |
| Surface righting | Pup placed supine on warm surface; released to flip over. | Latency to roll onto all fours (max 60 s). |
| Negative geotaxis | Pup placed head-down on 45° incline. | Latency to rotate 180° uphill (max 30 s). |
| Cliff aversion | Pup placed at edge of platform with forepaws over edge. | Latency to retreat (max 30 s). |
| Forelimb grasp | Pup positioned vertically so forepaws grasp rod. | Latency to fall during 30 s trial. |
| Rooting | Cheek stroked with cotton filament. | Head turns toward stimulus. |
| Ear twitch | Ear tip brushed gently with cotton filament. | Ear flattens in response. |
| Auditory startle | Handclap or finger snap near pup. | Quick startle movement observed. |

**Table S2A-J.**
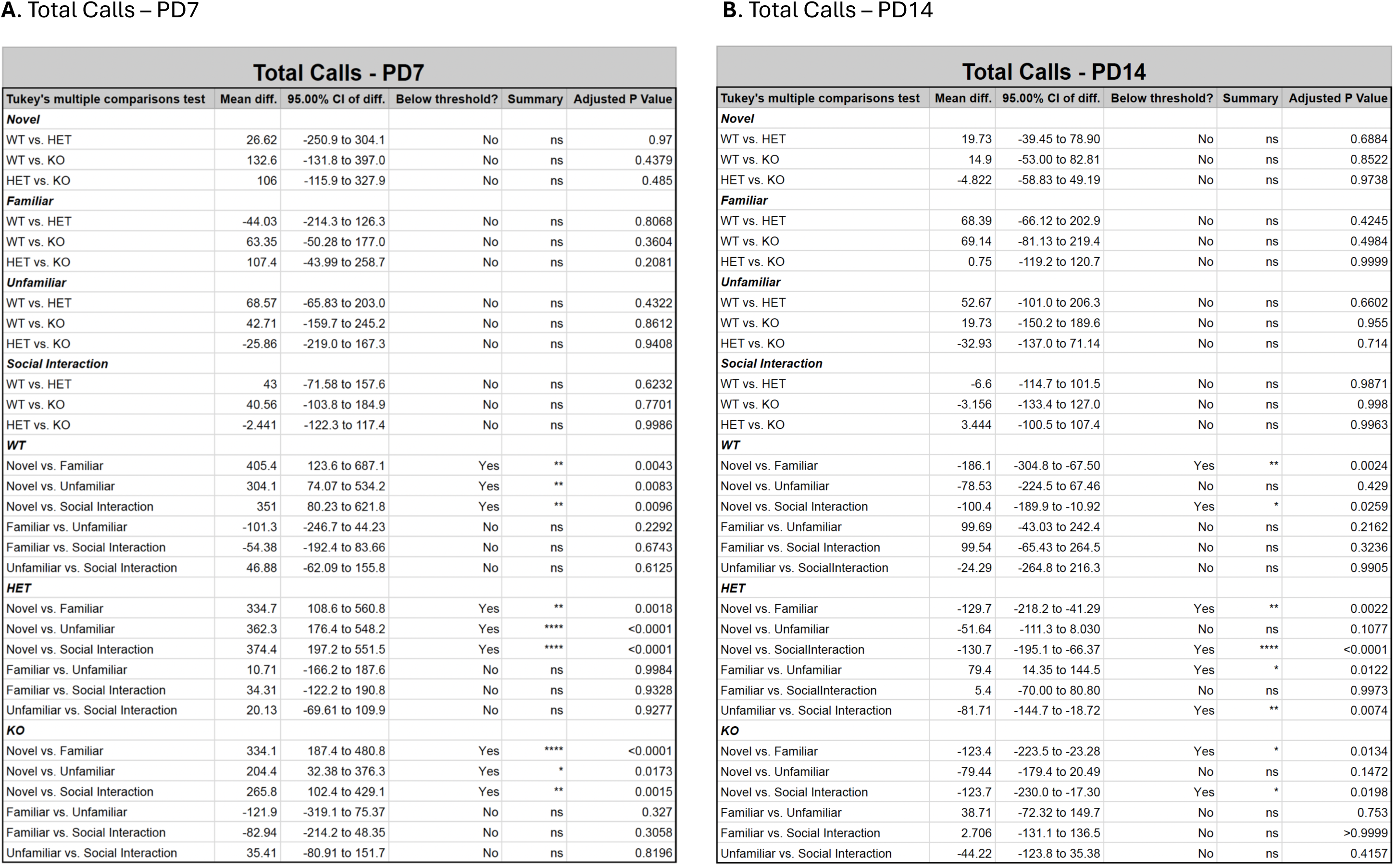
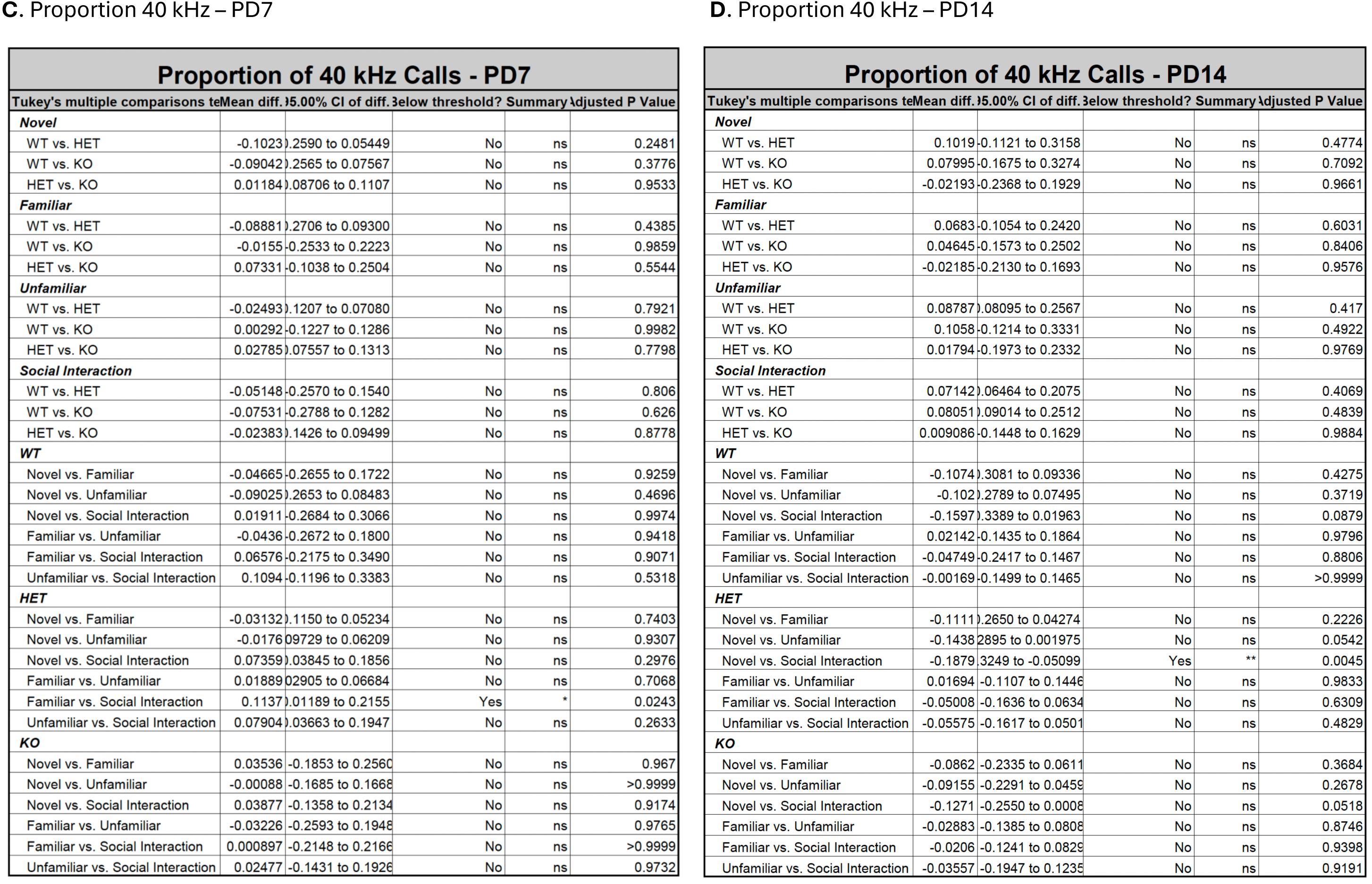
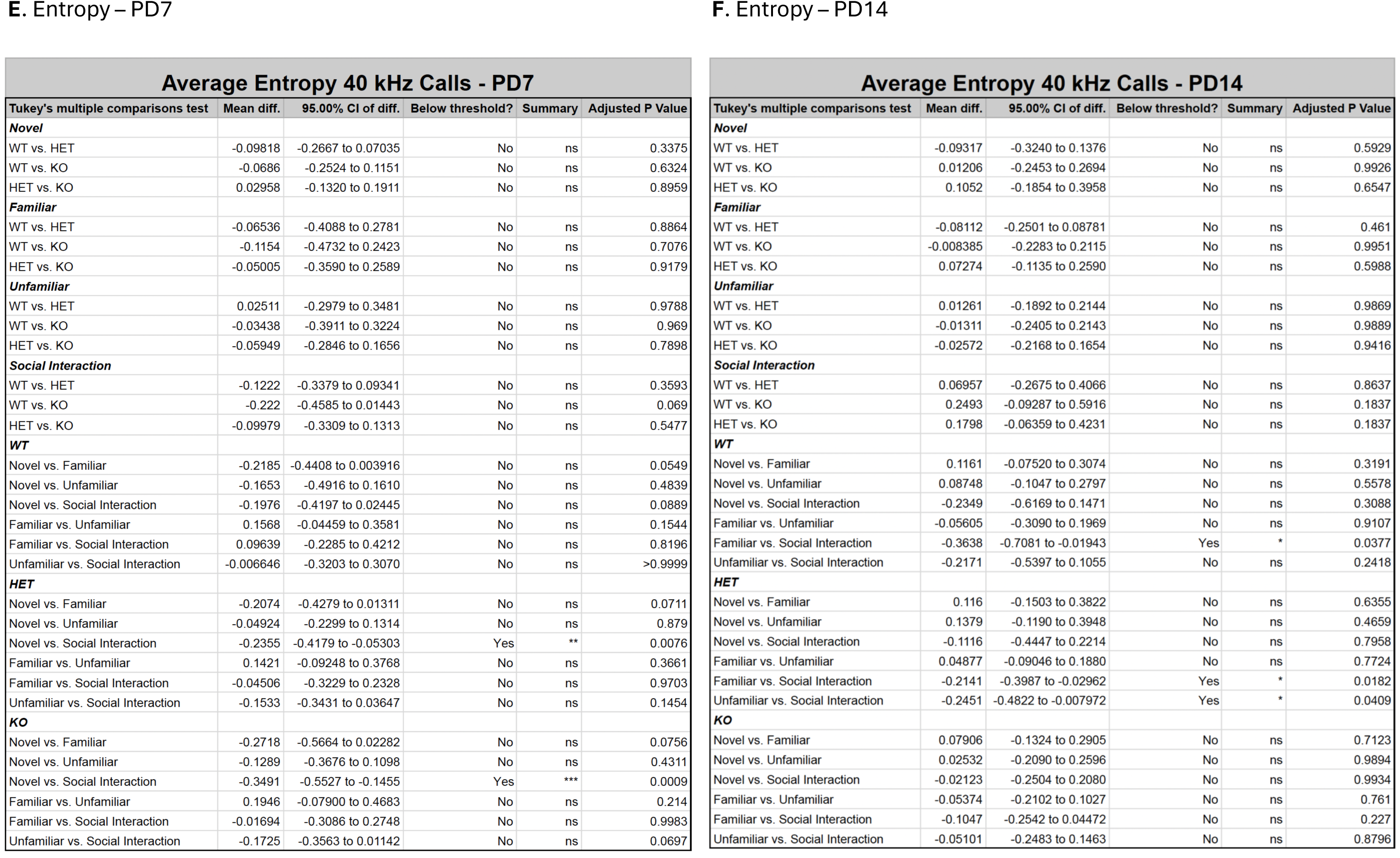
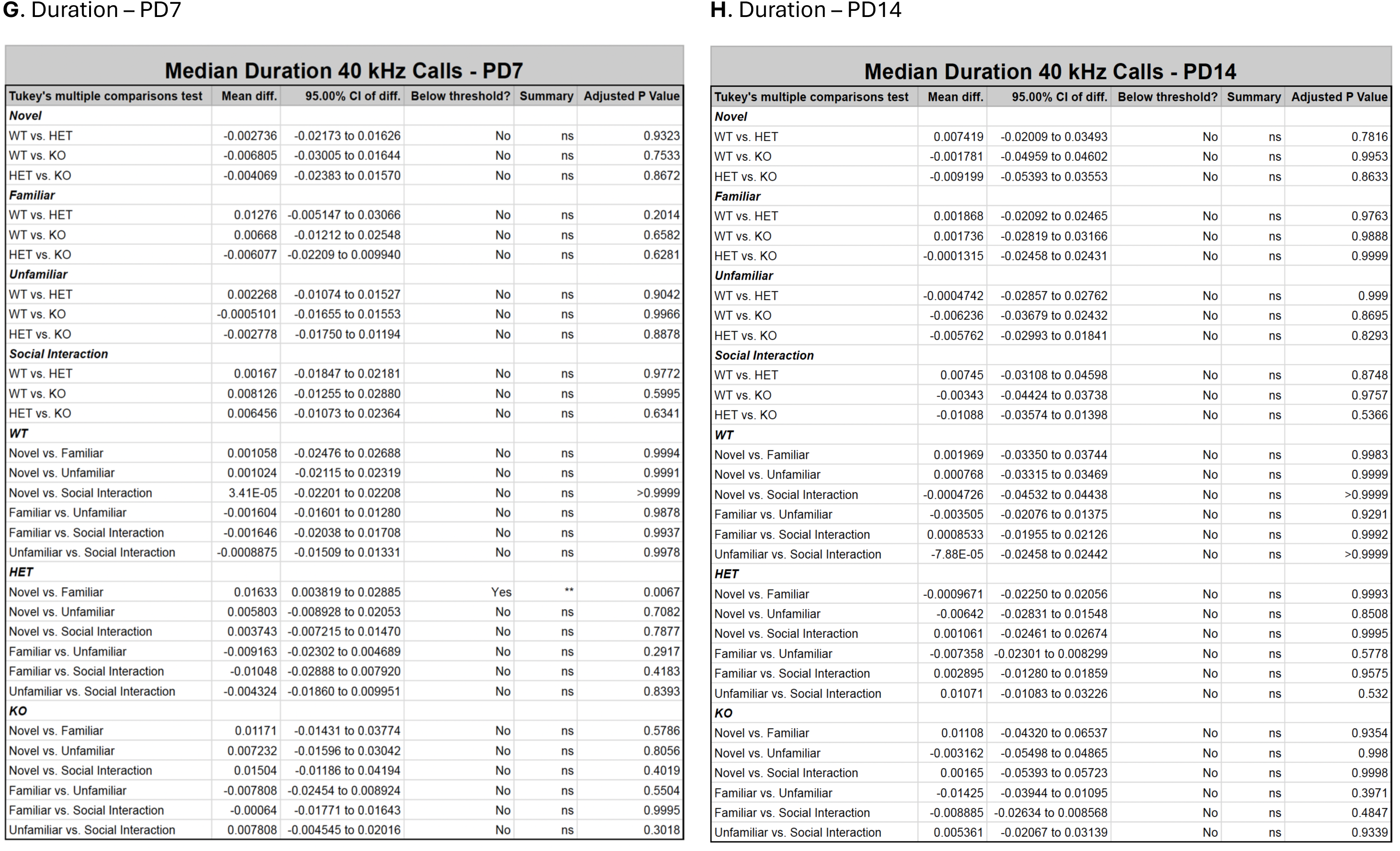
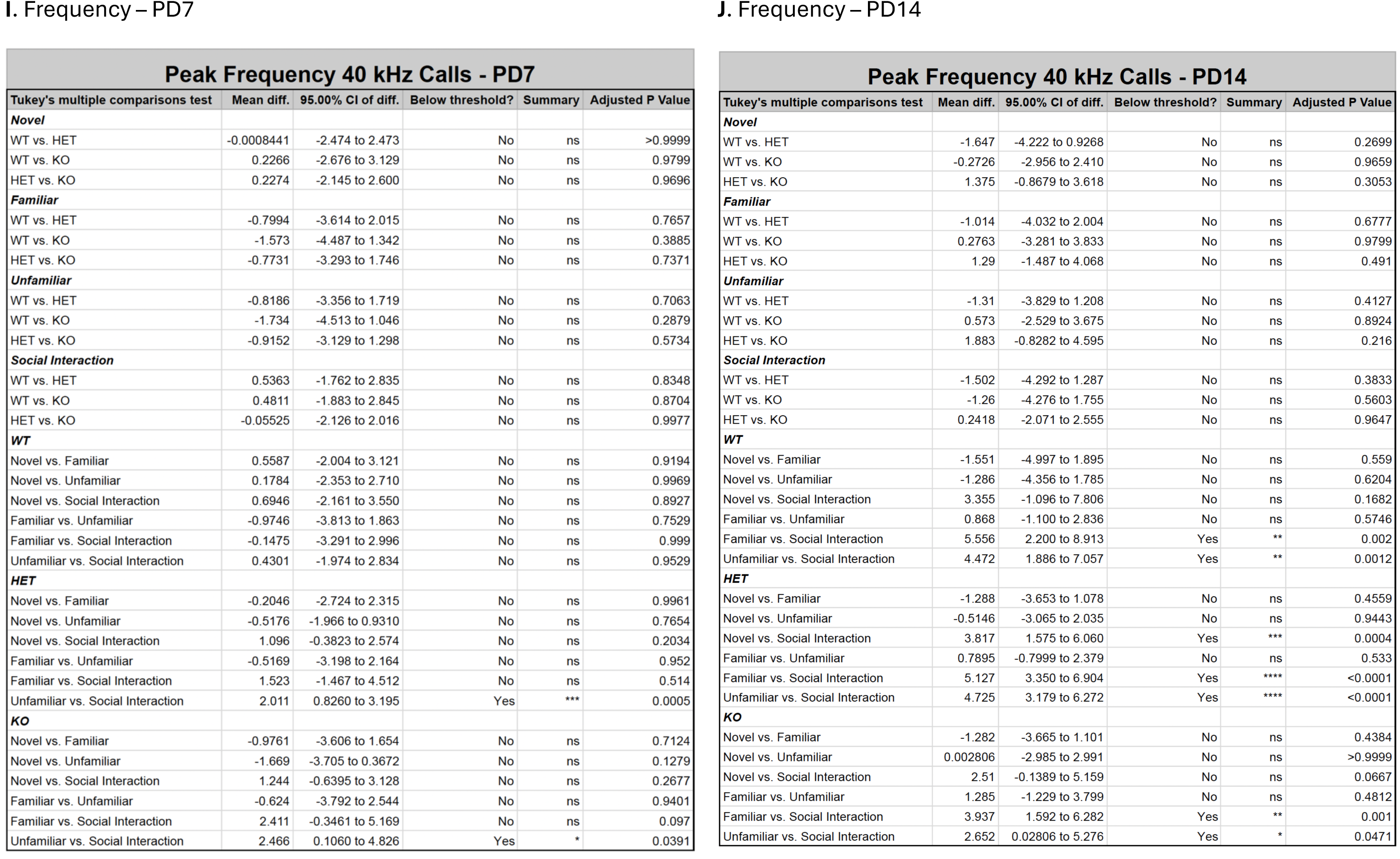
Statistical data for all group and condition comparisons for neonatal vocalizations.

**Figure. S1.**
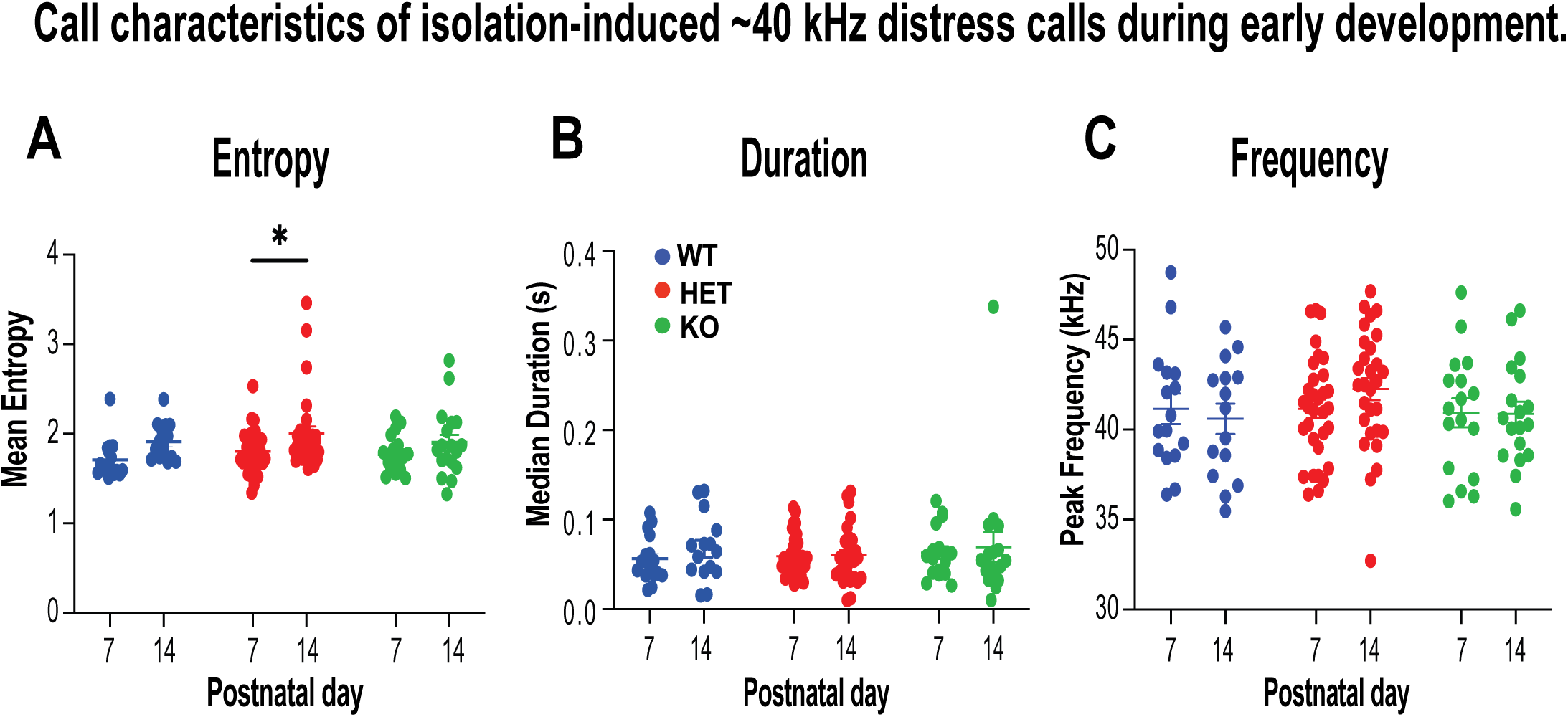
Call characteristics of isolation-induced ∼40 kHz distress calls during early development. **(A)**Mean call entropy of ∼40 kHz ultrasonic vocalizations recorded at postnatal day (PD) and PD14. The HET rats show an increase in entropy between PD7 and PD14, whereas WT and KO animals showed no age-dependent change. **(B)** Median call duration and **(C)** peak call frequency of ∼40 kHz vocalizations at PD7 and PD14 were not affected by genotype or age. Data are presented as mean ± SEM with individual animals shown as circles/dots. WT, wildtype; HET, heterozygous; KO, homozygous knockout. *P < 0.05.

**Figure. S2.**
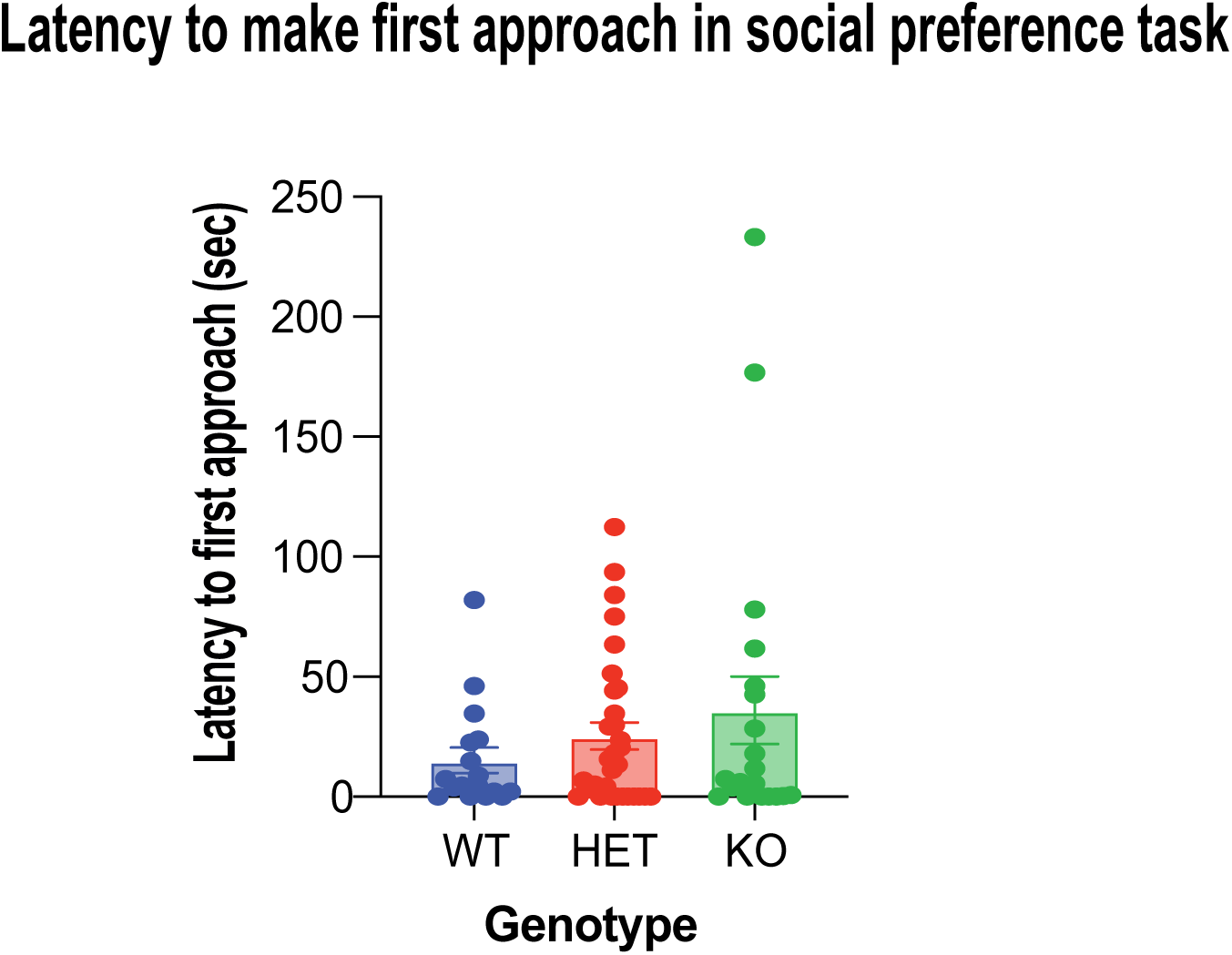
Latencies to make first social approach in juvenile Shank3-deficient rats. In social interaction task, latencies to approach and investigate preferred conspecific rat were comparable across groups. Thus, there was no evidence of impaired social approach behavior in juvenile Shank3-deficient rats.

**Figure. S3.**
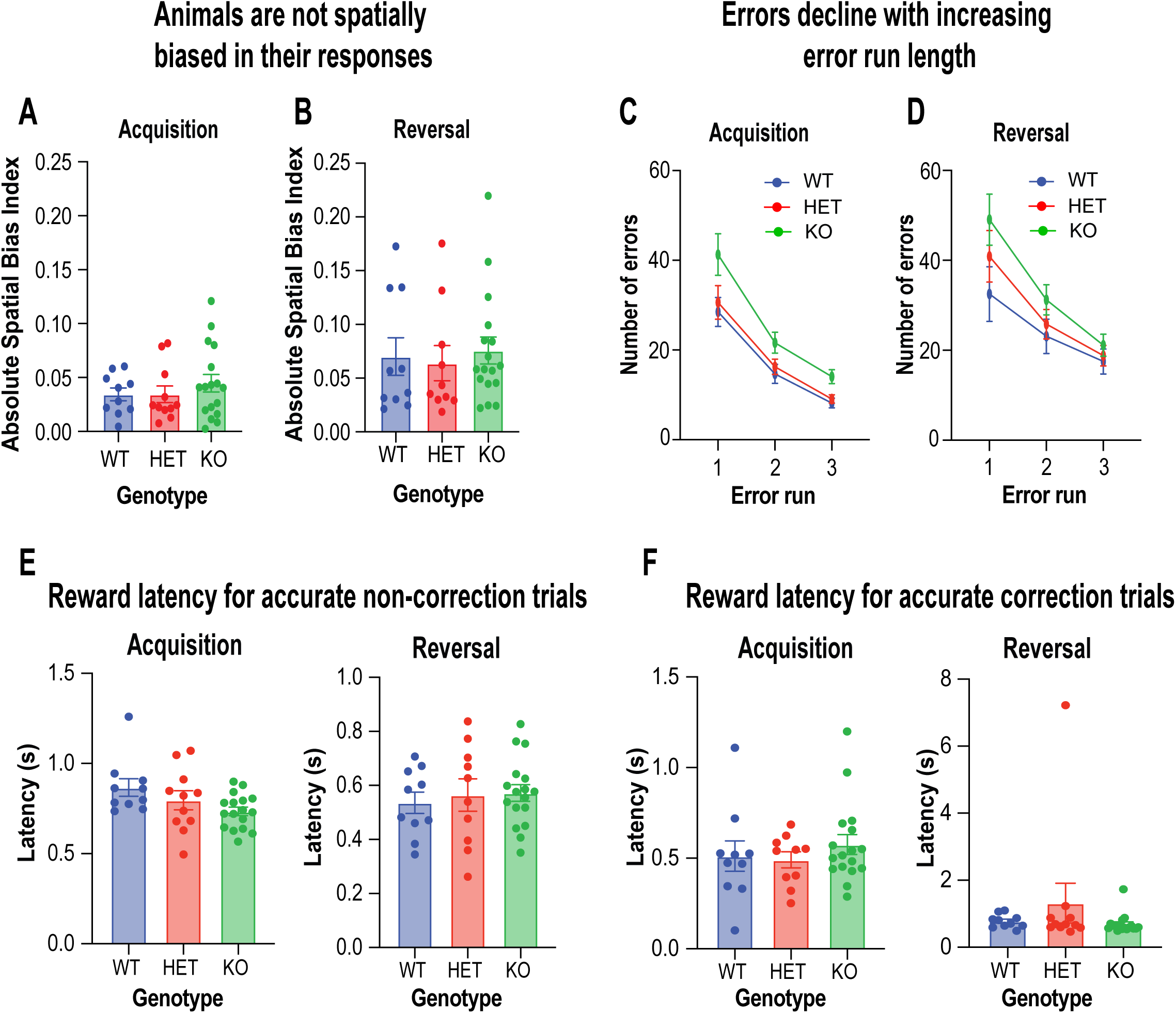
Touchscreen simple discrimination and reversal learning task in adult Shank3 mutant rats. Raw behavioral data were processed to compute a spatial bias index for each session, defined as (# left response trials − # right response trials) / total trials. This metric yields values ranging from−1 (complete rightward bias) to +1 (complete leftward bias), with 0 indicating no directional bias. For each animal, spatial bias was first quantified across sessions, and the absolute value of the index was averaged across trial types to assess overall bias magnitude independent of direction. This analysis revealed no significant differences in spatial bias between genotypes for the acquisition phase **(A)** nor the reversal phase **(B).** Errors committed during performance were further analyzed on a trial-by-trial basis to establish the likelihood of making an error following consecutive incorrect trials (error runs). On average, all animals corrected errors within 2–3 trials, making progressively few errors with increasing length of error run for both acquisition **(C)** and reversal **(D).** Thus, the correction trial errors exhibited by the KO rats were not driven by prolonged consecutive errors. Reward collection latencies for accurate responses for non-correction trials **(E)** and repeat correction trials (**F)** were largely unaffected by genotype.

